# Action of cocaine involves depletion of dopaminergic and serotonergic storage vesicles

**DOI:** 10.1101/651505

**Authors:** Judith R. Homberg, Peter Karel, Francisca Meyer, Kiki Rink, Josephus A. van Hulten, Nick H.M. van Bakel, Eric L.W. de Mulder, Lucia Caffino, Fabio Fumagalli, Jitske Jansen, Rosalinde Masereeuw, Gerard J.M. Martens, Alexander R. Cools, Michel M.M. Verheij

**Affiliations:** Department of Cognitive Neuroscience, division of Molecular Neurogenetics, Donders Institute for Brain, Cognition and Behaviour, Radboud University Nijmegen Medical Centre, Kapittelweg 29, 6525 EN Nijmegen, the Netherlands; Department of Molecular Animal Physiology, Radboud Institute for Molecular Life Sciences, Donders Center for Neuroscience, Faculty of Science, Radboud University Nijmegen, Geert Grooteplein Zuid 26-28, 6525 GA Nijmegen, The Netherlands; Department of Cognitive Neuroscience, division of Psychoneuropharmacology, Donders Institute for Brain, Cognition and Behaviour, Radboud University Nijmegen Medical Centre, Geert Grooteplein Noord 29, 6500 HB Nijmegen, the Netherlands; Department of Pharmacological and Biomolecular Sciences, Università degli Studi di Milano, Via Balzaretti 9, 20133 Milan, Italy; Division of Pharmacology, Utrecht Institute for Pharmaceutical Sciences, Utrecht University, Universiteitsweg 99, 3584 CG Utrecht, the Netherlands

**Keywords:** Cocaine pharmacology, cocaine mode of action, cocaine mechanism of action, vesicular monoamine release, storage vesicle depletion, synaptic vesicle depletion.

## Abstract

Cocaine is known to increase the extracellular levels of dopamine (DA) and serotonin (5-HT) by inhibiting the neuronal reuptake of these monoamines. However, individuals with reduced monoamine reuptake transporter expression do not display a reduction in cocaine intake, suggesting that a mechanism other than inhibition of monoamine reuptake contributes to the rewarding and addictive effects of the psychostimulant. Here we report that cocaine depletes the dopaminergic and serotonergic storage vesicles of the rat nucleus accumbens. This cocaine-induced vesicle depletion gave rise to acute increases in the extracellular levels of DA and 5-HT, which in turn correlated with monoamine-type-specific changes in behavior. Both the neurochemical and behavioral responses to cocaine varied among individual animals, which was not due to individual differences in the reuptake of DA and 5-HT, but rather to individual differences in their vesicular release. Furthermore, we found that reserpine-induced depletion of storage vesicles reduced both short and long access cocaine self-administration, and the degree of reduction was linked to the vesicular storage capacity of the animals. In conclusion, we demonstrate a novel mechanism by which cocaine increases the extracellular concentrations of accumbal DA and 5-HT, namely via release from storage vesicles. Furthermore, individual differences in cocaine-induced vesicular monoamine release shape individual differences in not only the acute behavioral and neurochemical effects of the stimulant, but also in its intake. Thus, intracellular storage vesicles represent an attractive novel drug target to combat psychostimulant addiction.

## Introduction

Cocaine (COC) is a powerful psychostimulant that produces short-term euphoria and often leads to addiction. Addiction to COC not only destroys lives of users, but also of their relatives. Presently, there are no FDA-approved drugs to treat COC addiction. Dopamine (DA) is well known to mediate COC effects [1–4], but recent evidence suggests that serotonin (5-HT) plays an important role in the neurochemical and behavioral effects of COC as well [5–9]. For instance, COC administration increases whereas COC withdrawal decreases the extracellular levels of both DA and 5-HT in the nucleus accumbens [9–13]. Moreover, in addition to accumbal DA receptors, accumbal 5-HT receptors have been found to mediate COC-induced locomotor activity and COC reward [14–20].

COC is known to increase extracellular monoamine levels by blocking plasmalemmal monoamine reuptake transporters [21]. However, the behavioral response to and the intake of COC is not reduced in animals and humans that are marked by a reduction of monoamine reuptake transporters [7, 8, 22–26]. In addition, we and others have found that an acute COC challenge still resulted in an increase in the extracellular levels of DA and 5-HT in the nucleus accumbens of animals lacking plasmalemmal DA and 5-HT transporters [9, 27–29]. False neurotransmitter uptake could explain these findings, but it has been shown that COC-induced increases in the accumbal extracellular levels of a particular monoamine (eg DA) are not explained by COC-mediated inhibition of another monoamine (eg 5-HT) transporter [30]. These findings suggest that a mechanism other than DA and 5-HT reuptake inhibition may contribute to the rewarding and addictive effects of COC. Interestingly, in 1977 Scheel-Krüger and co-workers have shown that the behavioral response to COC also depends on the presence of monoaminergic storage vesicles [see 31]. These observations led us to the hypothesis that COC releases DA and 5-HT into the synaptic cleft by depleting presynaptic storage vesicles.

In the present study, we have isolated the presynaptic monoaminergic storage vesicles of the rat nucleus accumbens, and measured COC-induced changes in the intra-vesicular as well as extracellular levels of accumbal DA and 5-HT. We show that COC depletes accumbal dopaminergic and serotonergic storage vesicles. Furthermore, we demonstrate that this COC-induced release of vesicular DA and 5-HT correlates with an increase in walking and rearing, respectively, and that individual-specific increases in the extracellular DA and 5-HT response to COC are due to individual differences in vesicular monoaminergic storage capacity. Finally, we show that depletion of monoaminergic storage vesicles inhibits the voluntary short and long access intake of COC.

## Materials and methods

### Rats

All experiments were performed in adult male Wistar rats, bred at the Central Animal Facility of the Radboud University, Nijmegen, the Netherlands. Given the individual differences in COC-induced vesicle depletion observed during the first experiment, the remaining 3 experiments were performed in High Responder (HR) and Low Responder (LR) to novelty rats, which have previously been found to differ in the expression of vesicular monoamine transporters type 2 (VMAT-2) [32]. All experiments were performed in accordance with institutional, national and international laws and guidelines for animal care and welfare (see National Research Council (NRC) 2003 guidelines). Every effort was made to minimize the number of animals used and their suffering.

### Experiment 1

Unselected rats received a single injection of COC (dose: 45 mg/kg) or saline (volume: 1 ml/kg, i.p.) and walking and rearing behavior was recorded. After 20 min, these rats were sacrificed by decapitation, bilateral punches of nucleus accumbens sections were collected [33, 34], and storage vesicles were isolated by ultracentrifugation according to the protocol of Staal *et al*., 2000 [35]. Afterwards, the accumbal vesicles were disrupted by osmotic shock and sonication and their content was injected into a High Performance Liquid Chromatography (HPLC) system coupled to an electrochemical detector (ECD) for the separation and quantification of the vesicular levels of DA and 5-HT.

### Experiments 2-4

HR and LR to novelty rats were selected on a novel open field according to procedures previously described by Cools *et al*., 1990 [36] and the accumbal levels of plasmalemmal DA and 5-HT uptake [for procedures, see 37, 38], in addition to the accumbal total and vesicular levels of DA and 5-HT (for procedures, see above), were measured. A second group of HR and LR rats underwent stereotactic surgery to implant a microdialysis probe into the right nucleus accumbens [see 39]. After recovery, these rats were injected with the vesicle depleting agent reserpine (RES, doses: 1 or 2 mg/kg) or its solvent (volume: 1 ml/kg, i.p.), and 24 h later accumbal levels of extracellular DA and 5-HT were measured using the above-mentioned HPLC-ECD setup. When the RES-induced reduction of the extracellular levels of accumbal DA and 5-HT was stable, the HR and LR rats of the second group were also injected with COC (dose: 15 mg/kg) or saline (volume: 1 ml/kg, i.p.), where after the extracellular accumbal levels of these monoamines as well as walking and rearing were measured for an additional period of 120 min. A third, and final, group of HR and LR rats was implanted with a catheter into the right external jugular vein [40] and, after training, COC self-administration (0.5 mg/kg/infusion) was performed for daily 1-h (short access) or 6-h (long access) sessions [for details 41, 42]. When the COC intake reached a stable level, rats were also treated with RES (dose: 1 or 2 mg/kg) or its solvent (volume: 1 ml/kg, i.p.), where after they were exposed to their corresponding COC self-administration sessions for an additional period of 6 days.

A detailed version of the experimental procedures and the subjects used can be found in the Supplemental Materials and Methods.

## Results

### COC results in vesicular monoamine release that is correlated with changes in walking and rearing behavior

Ultracentrifugation was used to isolate the storage vesicles of the rat nucleus accumbens (see Fig. 1a for location punch needle). These vesicles expressed high levels of various glycosylated forms of VMAT-2 (Fig. 1b). Sonication-induced disruption of these vesicles resulted in the liberation of vesicular DA (one-way ANOVA: sonication effect: (Fig. 1c): F_(1,14)_=5.329, p=0.037) and 5-HT (one-way ANOVA: sonication effect: (Fig. 1d): F_(1,14)_=13.187, p=0.003).

**Fig. 1:**
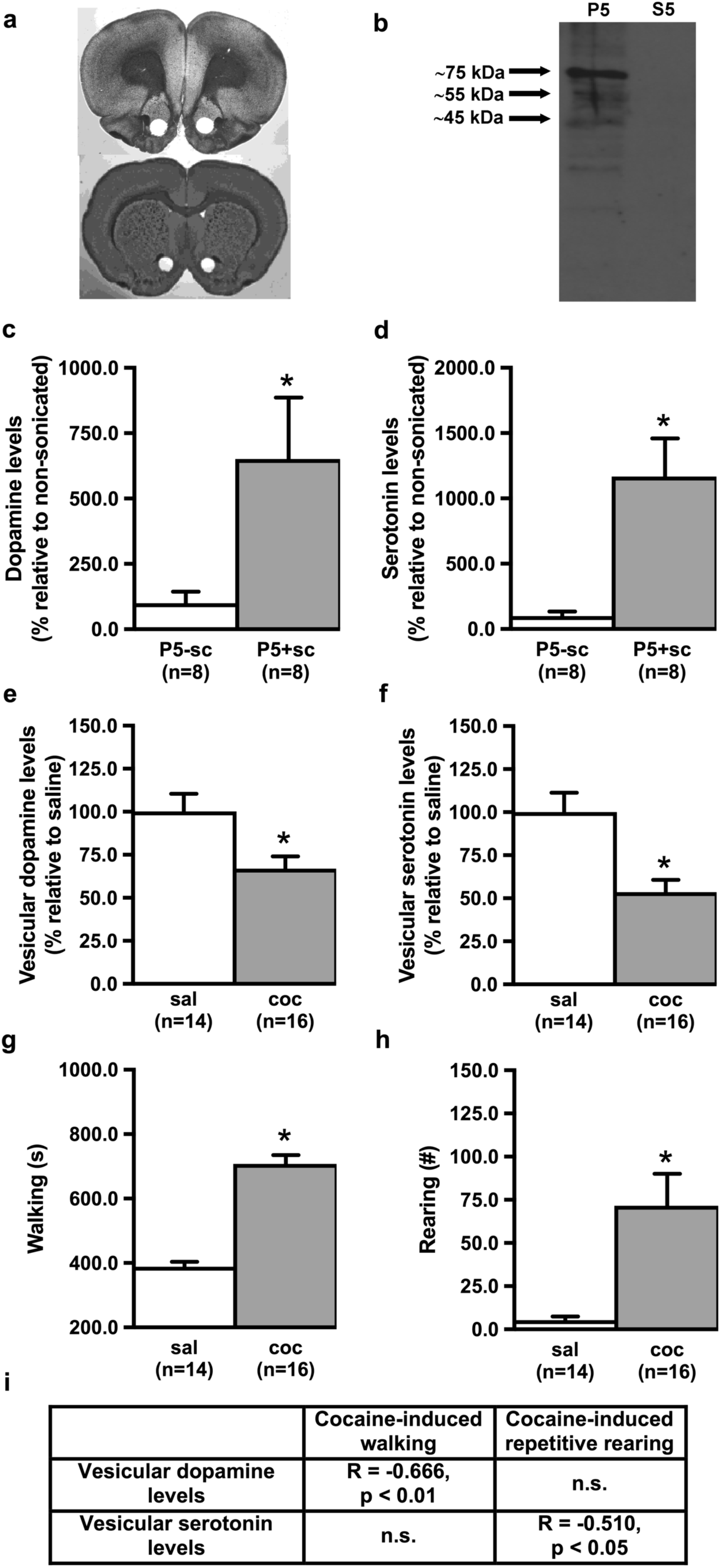
Cocaine (COC) results in vesicular monoamine release that is correlated with changes in walking and rearing behavior. Ultracentrifugation was used to isolate the monoaminergic storage vesicles of the rat nucleus accumbens (**a)** and Western blot analysis revealed various glycosylated forms of vesicular monoamine transporters type 2 (VMAT-2) in vesicular pellet P5, but not in its supernatant S5 (**b**). The vesicular pellet P5 and its supernatant S5 resulted from the ultracentrifugation of supernatant S4 at 100000g for 45 min (for details: see Supplemental Materials and Methods). Sonication (sc) of the re-suspended pellet P5 leads to the liberation of dopamine (DA) (**c**) and serotonin (5-HT) (**d**) from the isolated storage vesicles, and systemic administration of COC depletes these vesicles as demonstrated by a COC-induced reduction in the intra-vesicular levels of DA (**e**) and 5-HT (**f**). Systemic administration of COC also changed walking (**g**) and rearing (**h**). The COC-induced depletion of the dopaminergic storage vesicles of the nucleus accumbens significantly correlated with an increase in COC-induced walking, but not rearing (**i**). In contrast, the COC-induced depletion of the serotonergic storage vesicles of the nucleus accumbens significantly correlated with an increase in COC-induced rearing, but not walking (**i**). *Significant difference between sonicated (+sc) and non-sonicated (-sc) P5 vesicular fraction or between saline (sal)- and COC-treated rats (one-way ANOVA: p<0.05). All data are expressed as mean ± SEM.

If COC, as suggested in the introduction, releases monoamines from storage vesicles, it is expected that, after COC treatment, DA and 5-HT concentrations inside these vesicles will be reduced. Following a single systemic injection of COC, the amount of DA inside the storage vesicles of the rat nucleus accumbens was indeed reduced by ∼35% (one-way ANOVA: treatment effect (Fig. 1e): F_(1,28)_=7.296, p=0.012). The COC-induced depletion of the serotonergic storage vesicles in the nucleus accumbens was even larger (∼55%; one-way ANOVA: treatment effect (Fig. 1f): F_(1,28)_=12.695, p=0.001).

COC increased both walking (one-way ANOVA: treatment effect (Fig. 1g): F_(1,28)_=96.645, p<0.001) and rearing (one-way ANOVA: treatment effect (Fig. 1h): F_(1,28)_=11.834, p=0.002). COC-induced changes in behavior as well as vesicular monoamine depletion were higly individual-specific (Supplemental Fig. S1). Interestingly, the COC-induced depletion of vesicular DA (Fig. 1e) significantly correlated with the COC-induced increase in walking (Fig. 1i), but not rearing (Fig. 1i). In contrast, the COC-induced depletion of vesicular 5-HT (Fig. 1f) significantly correlated with the COC-induced increase in rearing (Fig. 1i), but not walking (Fig. 1i).

### Individual differences in vesicular DA and 5-HT levels

We have previously shown that rats with a high locomotor response (HR) to novelty express more VMAT-2 in their nucleus accumbens when compared to rats with a low locomotor response (LR) to novelty [32]. VMAT-2 is responsible for the transport of both DA and 5-HT into vesicles [43]. We therefore investigated whether animals that differ in their locomotor response to novelty also differ in their dopaminergic and serotonergic storage capacity of the nucleus accumbens.

HR and LR rats were selected on the basis of their locomotor response to a novel open-field (Supplemental Fig. S2a). The average distance travelled in 30 min was 3.493 ± 191 cm for LR and 8.643 ± 373 cm for HR. The average habituation time (for details, see supplemental Materials and Methods) was 324 ± 28 s and 1.340 ± 66 s for LR and HR, respectively. The amount of both DA and 5-HT stored inside accumbal storage vesicles was larger in HR than in LR (one-way ANOVA: rat type effect: DA (Supplemental Fig. S2b): F_(1,16)_=6.100, p=0.025 and 5-HT (Supplemental Fig. S2c): F_(1,16)_=8.150, p=0.011). These higher levels of vesicular DA and 5-HT in HR than in LR may account for the finding that the total accumbal levels of DA and 5-HT were also higher in HR as compared to LR (one-way ANOVA: rat type effect: DA (Supplemental Fig. S2d): F_(1,18)_=6.000, p=0.023 and 5-HT (Supplemental Fig. S2e): F_(1,18)_=5.606, p=0.029). A Pearson’s analysis revealed that both the travelled distance and the habituation time, assessed during the open-field selection procedure, positively correlated with the vesicular levels of accumbal DA and 5-HT (Supplemental Fig. S2f), as well as the total accumbal levels of these monoamines (Supplemental Fig. S2f).

### Individual differences in DA and 5-HT response to COC

The finding that the amount of DA and 5-HT inside the storage vesicles of the nucleus accumbens are larger in HR than in LR rats (Supplemental Fig. S2), together with the finding that COC releases monoamines from these storage vesicles (Fig. 1), suggests that COC increases the extracellular levels of these monoamines more strongly in HR than in LR rats. We found that 15 mg/kg COC increased the extracellular accumbal DA and 5-HT levels in HR (three-way ANOVA: treatment x time effect: DA: F_(28,420)_=6.304, p≤0.001 and 5-HT: F_(28,420)_=35.875, p≤0.001) and in LR (three-way ANOVA: treatment x time effect: DA: F_(28,392)_=3.381, p≤0.001 and 5-HT: F_(28,392)_=8.923, p≤0.001). However, the COC-induced increase in these extracellular monoamine levels was significantly larger in HR than in LR (three-way ANOVA: rat type x treatment x time effect: DA (Fig. 2a): F_(28,812)_=1.855, p=0.005 and 5-HT (Fig. 2b): F_(28,812)_=7.069, p≤0.001). Both the travelled distance and the habituation time, assessed during the open-field selection procedure, positively correlated with the maximum COC-induced increase in accumbal DA and 5-HT (Fig. 2c).

**Fig. 2:**
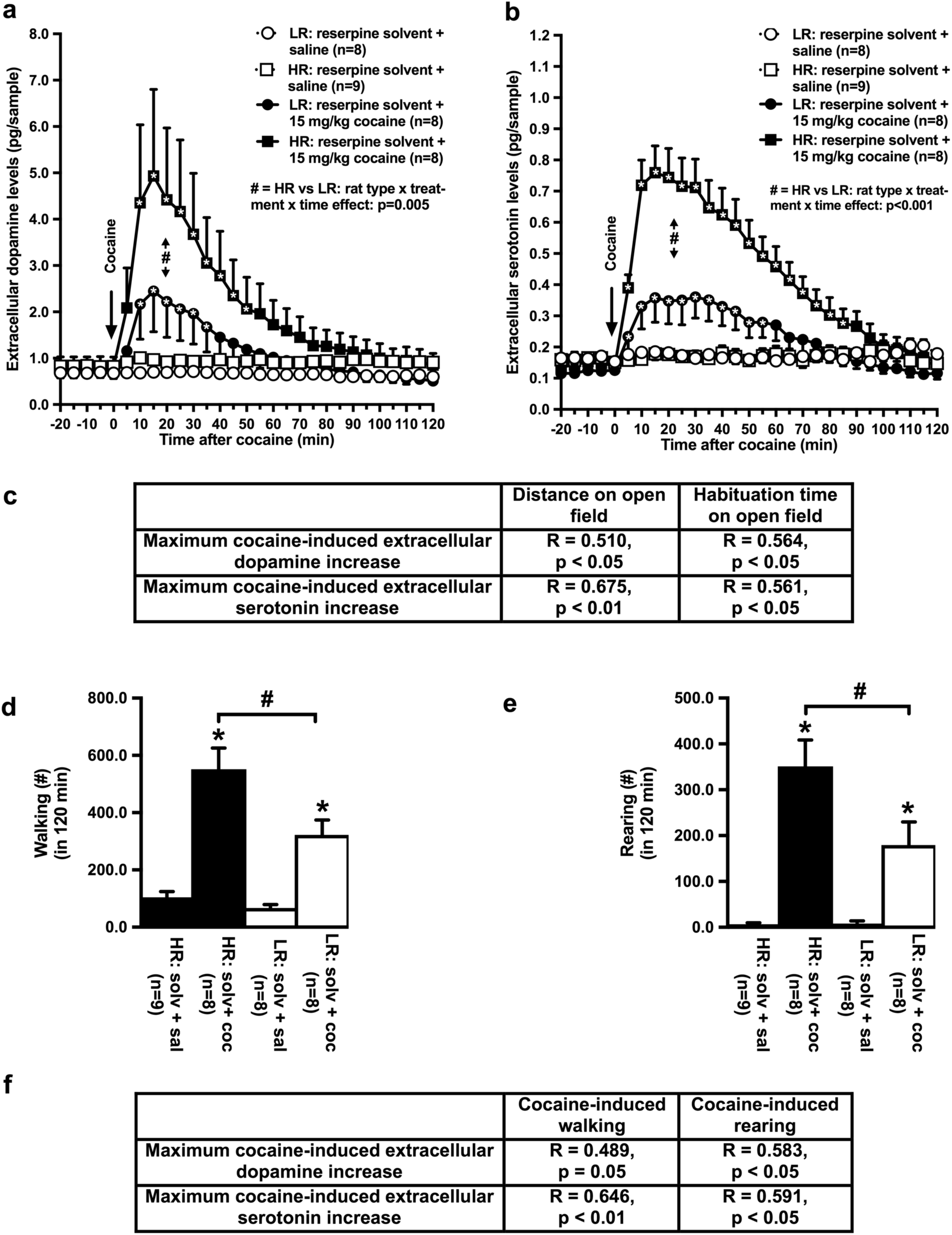
Individual differences in monoamine and behavioral responses to cocaine (COC). Systemic administration of COC increases the extracellular levels of accumbal dopamine (DA) (**a)** and serotonin (5-HT) (**b**) more strongly in High Responder to novelty (HR) than in Low Responder to novelty (LR) rats, and the maximum COC-induced increase in the extracellular levels of these monoamines was found to correlate significantly with the distance travelled and habituation time used to select these animals on the open-field (**c**). 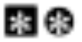 Significant COC-induced monoamine increase relative to saline (Student’s t-test: p<0.05). ^#^Significant difference between HR and LR (multi-way ANOVA: p≤0.005). COC also increased the frequency of walking (**d**) and rearing (**e**) more strongly in HR than in LR. The observed COC-induced increase in the extracellular levels of DA and 5-HT correlated with the increase in COC-induced walking and rearing (**f**). *Significant difference between COC and saline (sal), and ^#^significant difference between HR and LR (one-way ANOVA: p<0.05). Correlation analysis was performed in the pooled group of HR and LR. All rats were pre-treated with the solvent (solv) or reserpine (RES). All data are expressed as mean ± SEM.

### Individual differences in walking and rearing response to COC

Our HR and LR rats have previously been shown not to differ in the *duration* of walking and rearing following the administration of 15 mg/kg COC [44]. This dose of COC, however, is known to increase the number of automatically collected locomotor activity *counts* more strongly in HR than in LR [45]. Here we found that 15 mg/kg COC indeed increases the *frequency* of walking (two-way ANOVA: treatment effect: F_(1,29)_=61.584, p≤0.001) and rearing (two-way ANOVA: treatment effect: F_(1,29)_=48.110, p≤0.001) events more strongly in HR than in LR (two-way ANOVA: rat type x treatment effect: walking (Fig. 2d) F_(1,29)_=4.626, p=0.040 and rearing (Fig. 2e): F_(1,29)_=5.491, p=0.026). The COC-induced increase in walking and rearing correlated not only with the COC-induced reduction in the intra-vesicular levels of DA and 5-HT (see Fig. 1), but also with the COC-induced increase in the extracellular levels of these monoamines (Fig. 2f).

### Monoaminergic storage vesicles control individual differences in DA and 5-HT response to COC

To study the putative contribution of storage vesicles to the observed individual differences in the COC-induced increase in extracellular DA and 5-HT levels, COC-treated HR and LR rats were pre-treated with the VMAT-2 inhibitor reserpine (RES). After RES treatment, storage vesicles are known to become empty [39, 46–49]. Because accumbal storage vesicles contain more monoamines in HR than in LR (Supplemental Fig. S2), we hypothesized that COC-treated HR are less sensitive to the monoamine-depleting effects of RES than COC-treated LR.

1 mg/kg RES reduced baseline levels (Supplemental Fig. S3) and the COC-induced increase in the extracellular levels of DA and 5-HT (three-way ANOVA: treatment x time effect: DA (Figs. 3a and 3c): F_(28,784)_=3.127, p≤0.001 and 5-HT (Figs. 3b and 3d): F_(28,784)_=8.934, p≤0.001) in both types of rats (three-way ANOVA: rat type x treatment (x time) effect: DA and 5-HT: n.s.). However, the finding that COC could still increase accumbal extracellular DA and 5-HT levels in HR treated with 1 mg/kg RES (Figs. 3a and 3b: one sample t-tests vs baseline: P<0.05), but not anymore in LR treated with this dose of the vesicle depleting agent (Figs. 3c and 3d: one sample t-tests vs baseline n.s.) confirms our hypothesis that COC-treated HR are less sensitive to RES than COC-treated LR.

**Fig. 3:**
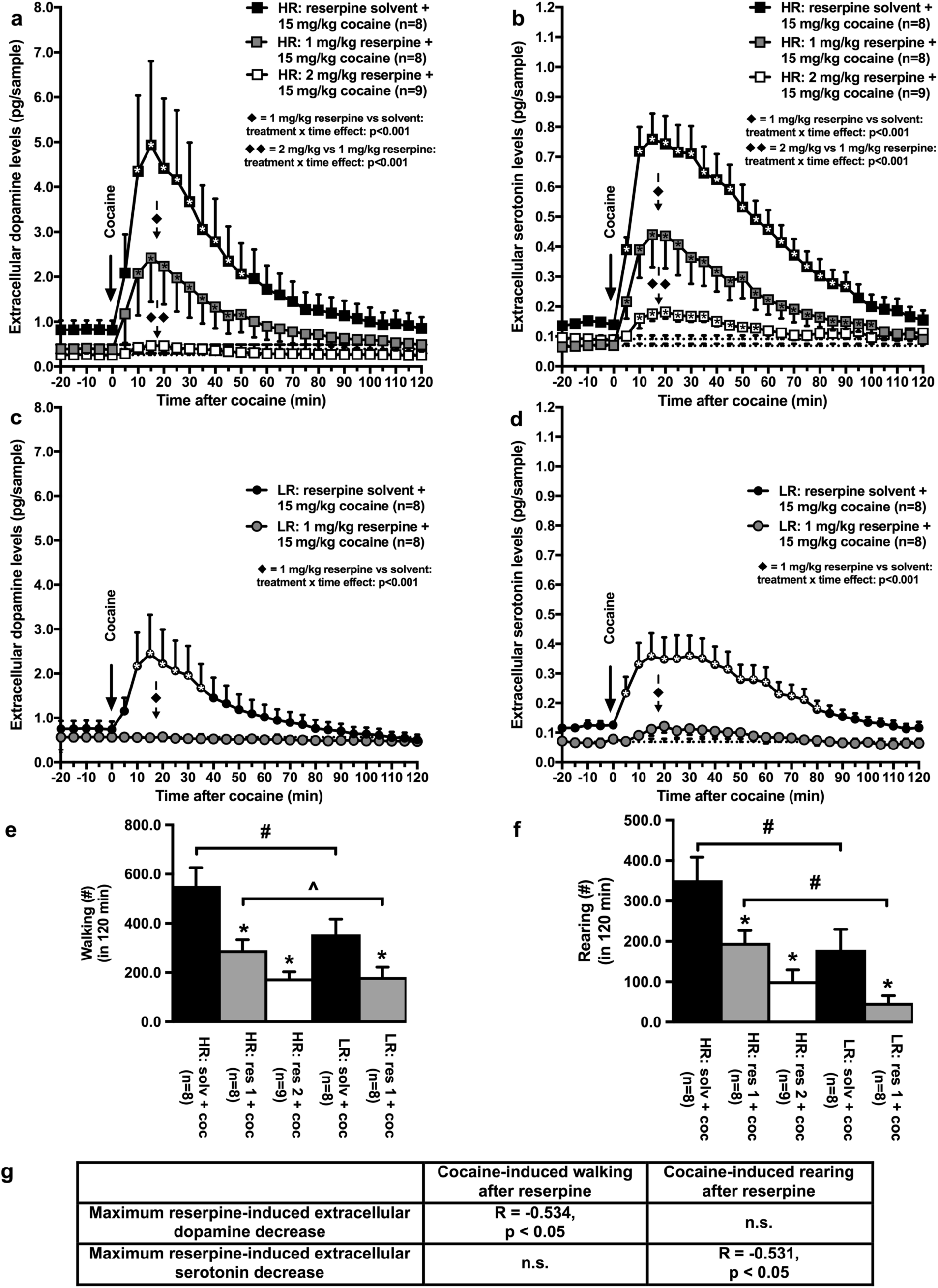
Monoaminergic storage vesicles control individual differences in monoamine and behavioral responses to cocaine (COC). Reserpine (RES) reduced the COC-induced increase in accumbal extracellular dopamine (DA) and serotonin (5-HT) levels in both High Responder to novelty (HR) (**a/b**) and Low Responder to novelty (LR) (**c/d**) rats. However, COC could still increase the accumbal extracellular monoamine levels in HR, but not anymore in LR, that were treated with 1 mg/kg of the vesicle depleting agent. 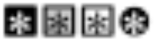 Significant COC-induced monoamine increase relative to baseline (one-sample t-tests: p<0.05). ♦ Significant RES-induced monoamine reduction, and ♦♦ significant RES dose effect (multi-way ANOVA: p<0.05). RES also reduced the COC-induced increase in walking and rearing in HR and in LR (**e/f**). *Significant decrease after RES (one-way ANOVA: p<0.05). ^#^Significant difference between HR and LR (one-way ANOVA: p<0.05). ^Trend towards significant difference between HR and LR (one-way ANOVA: p<0.075). The observed RES-induced reduction in the extracellular levels of DA was significantly correlated with the decrease in walking, but not rearing (**g**), whereas the observed RES-induced reduction in the extracellular levels of 5-HT was significantly correlated with the decrease in rearing, but not walking (**g**). Correlation analysis was performed in the pooled group of HR and LR treated with 1 mg/kg of RES. All data are expressed as mean ± SEM.

To test whether the effects of RES on the COC-induced increase in the extracellular levels of accumbal DA and 5-HT were dose dependent, a new group of COC-treated HR was pre-treated with 2 mg/kg of RES. This higher dose of RES was not tested in LR, because 1 mg/kg of RES already completely prevented the COC-induced DA and 5-HT increase in these rats (Figs. 3c and 3d). As expected, 2 mg/kg RES reduced the COC-induced increase in extracellular DA and 5-HT (two-way ANOVA: treatment x time effect: DA: F_(28,420)_=5.907, p≤0.001 and 5-HT: F_(28,420)_=27.878, p≤0.001) more strongly than 1 mg/kg of the drug (two-way ANOVA: dose x time effect: DA (Fig. 3a): F_(28,420)_=3.976, p≤0.001 and 5-HT (Fig. 3b): F_(28,420)_=6.755, p≤0.001).

### Monoaminergic storage vesicles control individual differences in walking and rearing response to COC

The RES-induced reduction in the COC-mediated increase of accumbal DA and 5-HT (Figs. 3a-3d) prompted us to investigate vesicle-depletion-induced changes in COC-mediated behavior. 1 mg/kg RES reduced the frequency of COC-induced walking and rearing events (two-way ANOVA: treatment effect: walking (Fig. 3e): F_(1,28)_=14.818, p=0.001 and rearing (Fig. 3f): F_(1,28)_=11.513, p=0.002) in both HR and LR rats (two-way ANOVA: rat type x treatment effect: walking and rearing: n.s.). Interestingly, the RES-induced reduction in the COC-induced extracellular DA increase (Fig. 3a) correlated with a decrease in walking (Fig. 3g), but not rearing (Fig. 3g). In contrast, the RES-induced reduction in the COC-induced extracellular accumbal 5-HT increase (Fig. 3b) correlated with a decrease in rearing (Fig. 3g), but not walking (Fig. 3g).

The additional HR group of rats pre-treated with 2 mg/kg RES (see above) was also used to establish the putative dose-response relationship for the vesicle-depletion-induced regulation of COC-induced behavior. The dose of 2 mg/kg of RES reduced COC-induced walking and rearing (one-way ANOVA: treatment effect: walking: F_(1,15)_=27.059, p≤0.001 and rearing: F_(1,15)_=17.400, p=0.001) more strongly than 1 mg/kg of the drug (one-way ANOVA: dose effect: walking (Fig. 3e): F_(1,15)_=5.676, p=0.030 and rearing (Fig. 3f): F_(1,15)_=5.263, p=0.036).

### Reuptake of DA and 5-HT does not control individual differences in response to COC

The results reported above indicate that individual differences in the monoaminergic storage capacity of the nucleus accumbens contribute to individual differences in the behavioral and extracellular monoamine response to COC. Behavioral and neurochemical responses to COC have also been found to depend on plasmalemmal monoamine re-uptake transporters [50]. In addition, changes in neuronal monoamine re-uptake may contribute to changes in novelty-induced exploration [51]. Therefore, individual differences in the re-uptake of DA and 5-HT by the plasmalemmal dopamine or serotonin transporters (DAT or SERT) may also have contributed to the reported results. To explore this possibility, the neuronal DA and 5-HT uptake was investigated in synaptosomes isolated from the nucleus accumbens of HR and LR rats. We found that the neuronal DA and 5-HT uptake in the nucleus accumbens was concentration dependent (two-way ANOVA: concentration effect: DA: F_(5,105)_=118.100, p≤0.001 and 5-HT: F_(5,105)_=268.831, p≤0.001), but not different between HR and LR rats (two-way ANOVA: rat type (x concentration) effect: DA (Supplemental Fig. S4a): n.s. and 5-HT (Supplemental Fig. S4b): n.s.).

### Individual differences in COC self-administration

The individual-specific increase in the extracellular levels of accumbal DA and 5-HT observed after forced exposure to COC (Fig. 2) suggests that HR and LR also differ in the voluntary intake of the drug. To test this hypothesis, HR and LR rats were trained to self-administer COC, after which they were divided into 2 groups that had daily access to the psychostimulant during 1 or 6 hour(s) sessions [8, 40–42]. In contrast to short access (ShA: 1 hour per session) COC self-administration, long access (LgA: 6 hours per session) COC self-administration resulted in an escalation of the drug intake over time (three-way ANOVA: access x time effect (Fig. 4a): F_(14,224)_=3.543, p<0.001; time effect: LgA: F_(14,112)_=4.286, p<0.001 and ShA: n.s.). The LgA, but not ShA, COC intake was higher in HR than in LR rats (three-way ANOVA: rat type x access x time effect: n.s; rat type x access effect: F_(1,16)_=5.450, p<0.05; rat type effect: LgA: F_(1,8)_=5.391, p<0.05 and ShA: n.s.). In addition, COC intake correlated with the locomotor response to novelty (Fig. 4b).

**Fig. 4:**
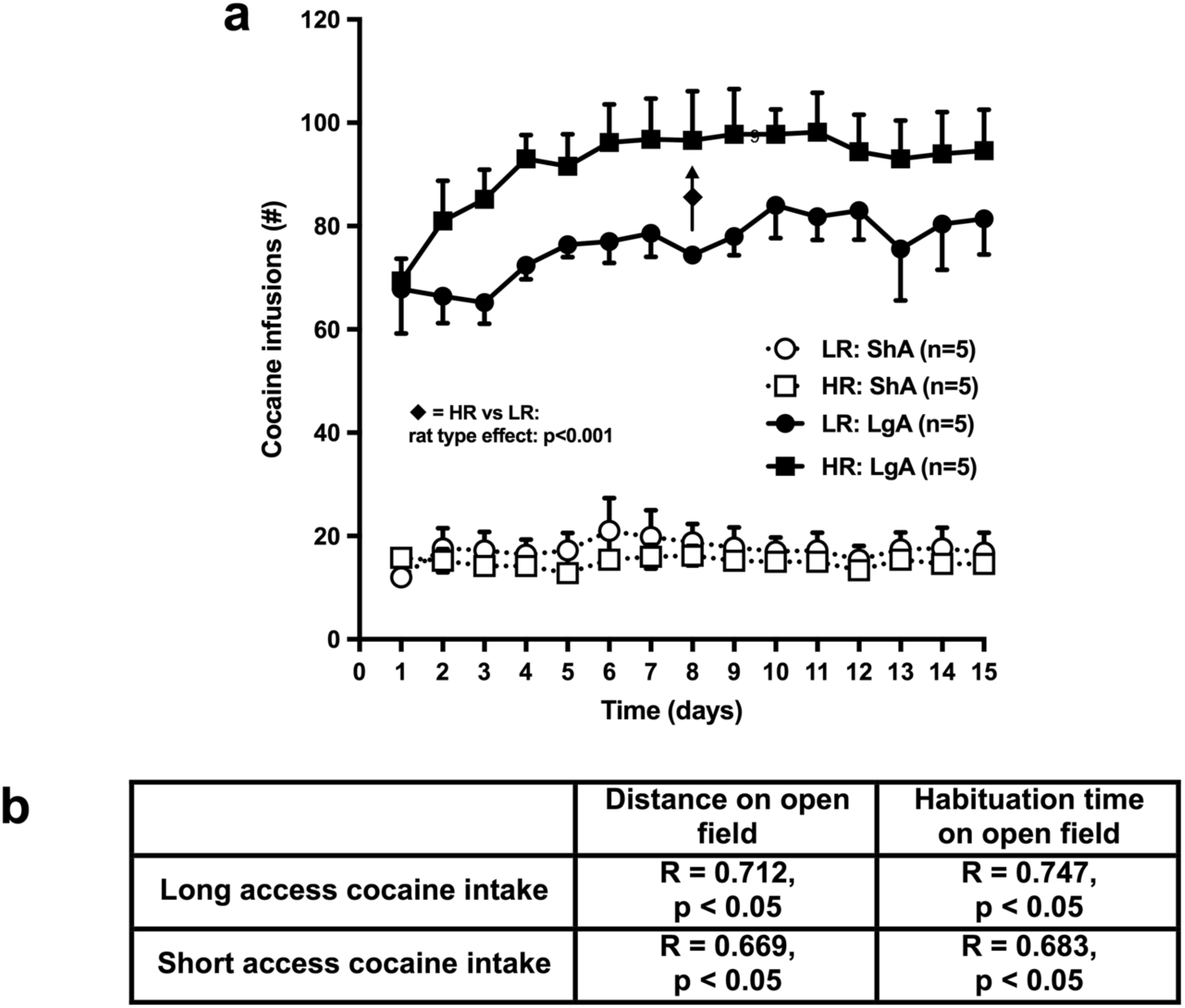
Individual differences in cocaine (COC) self-administration. Short Access (ShA) and Long Access (LgA) voluntary COC self-administration (**a**) correlated positively with the locomotor response to novelty (**b**). ♦ Significant rat type effect (multi-way ANOVA: p<0.05). Correlation analysis was performed in a pooled group of High Responder to novelty (HR) and Low Responder to novelty (LR) rats. All data are expressed as mean ± SEM.

### Monoaminergic storage vesicles control individual differences in COC self-administration

The observed RES-induced reduction in the COC-mediated increase of extracellular DA and 5-HT in the nucleus accumbens (Fig. 3) prompted us to investigate whether RES-induced depletion of monoaminergic storage vesicles also affects the COC intake in HR and LR rats. Since LR rats were marked by a smaller RES-sensitive monoaminergic storage capacity than HR rats (Supplemental Fig. S2), COC intake was hypothesized to be reduced by RES more strongly in LR than in HR rats. 1 mg/kg RES did not affect ShA COC self-administration (three way ANOVA: treatment (x time) effect (Figs. 5a and 5b): n.s.). However, the relatively low dose of RES reduced COC self-administration under LgA conditions (three-way ANOVA: treatment x time effect: n.s; treatment effect (Figs. 5c and 5d): F_(1,16)_=10.016, p<0.01). As hypothesized, LR rats were more sensitive to the RES-induced reduction in COC intake than HR rats (two-way ANOVA: rat type x treatment effect: ShA (Fig. 5e): n.s. and LgA (Fig. 5f): F_(1,16)_=5.248, p<0.05). In LgA rats, the maximum RES-induced decrease in COC intake (Fig. 5f) was negatively correlated with the locomotor response to novelty (Fig. 5g). No correlation was found between the locomotor response to novelty and the COC intake after applying 1 mg/kg RES to ShA rats (Fig. 5g), possibly because in these rats this dose of RES did not affect psychostimulant intake (Fig. 5e).

**Fig. 5:**
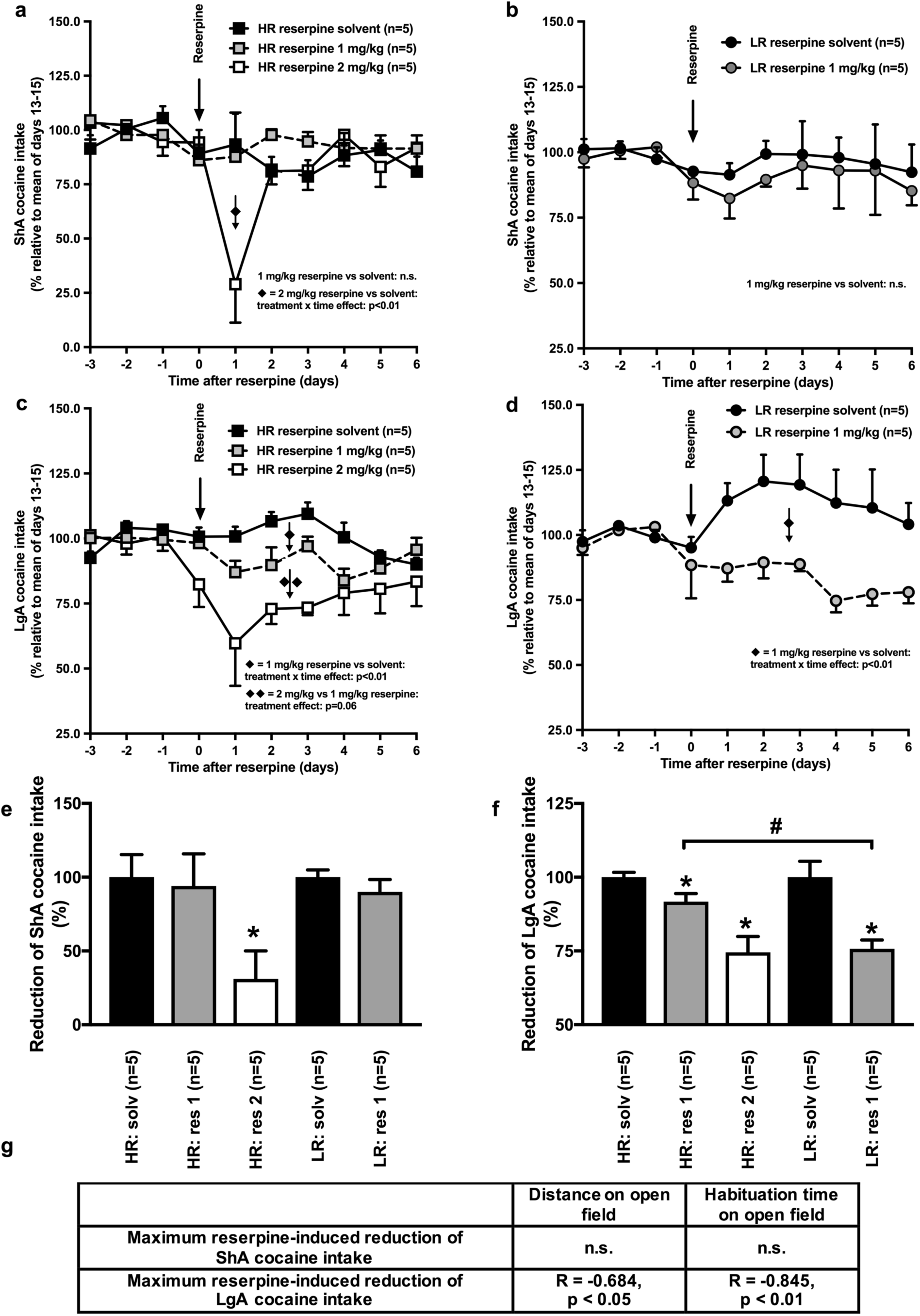
Monoaminergic storage vesicles control individual differences in cocaine (COC) self-administration. Reserpine (RES), 1 mg/kg, did not affect voluntary COC self-administration under Short Access (ShA) conditions (**a/b**). However, this low dose of the vesicle depleting agent reduced the Long Access (LgA) self-administation of the psychostimulant (**c/d**). ♦ Significant RES-induced reduction in the COC intake. The higher dose (2 mg/kg) of RES reduced not only the LgA, but also the ShA, COC intake (**a/c**). ♦♦ significant RES dose effect (multi-way ANOVA: p<0.05). The smaller RES-sensitive storage capacity in LR *versus* HR rats (see Supplemental Fig. S2) is in line with LR rats being more sensitive to RES-induced vesicle depletion than HR rats (**e/f**). *Significant decrease after RES (one-way ANOVA: p<0.05). ^#^Significant difference between HR and LR (one-way ANOVA: p<0.05). A low locomotor response to novelty correlated with a strong RES-induced reduction in the LgA COC intake (**g**). Correlation analysis was performed in the pooled group of HR and LR. Note: panels e and f represent the mean COC intake after RES treatment expressed as a percentage of the mean COC intake after treatment with RES solvent. 1 mg/kg of RES reduced the mean LgA intake of COC by 25% in LR and 10% in HR. All data are expressed as mean ± SEM.

To test whether the effects of RES on COC self-administration were dose dependent, a new group of COC-treated HR was pre-treated with 2 mg/kg of the vesicle depleting agent (see also Fig. 3). As expected, 2 mg/kg RES reduced the COC intake under both access conditions (two-way ANOVA: treatment x time effect: ShA (Fig. 5a): F_(9,72)_=3.218, p≤0.01 and LgA (Fig. 5c): F_(9,72)_=4.076, p≤0.001) more strongly than 1 mg/kg of the drug (one-way ANOVA: dose effect: ShA (Fig. 5e): F_(1,8)_=7.888, p≤0.05 and LgA (Fig. 5f): F_(1,8)_=8.066, p≤0.05).

## Discussion

### COC depletes monoaminergic storage vesicles

Our data confirm our hypothesis and demonstrate that systemic administration of COC depletes VMAT-2-containing dopaminergic and serotonergic storage vesicles in the nucleus accumbens of the rat. The finding that RES-induced depletion of these storage vesicles prior to the administration of COC significantly reduced the COC-induced increases in the extracellular levels of accumbal DA and 5-HT, as well as COC-mediated walking and rearing, shows that COC-induced vesicle depletion results in an increase of neurotransmitters in the synaptic cleft. Under the condition that COC depletes presynaptic storage vesicles, replenishment of these vesicles is required. Indeed, enhanced VMAT-2-dependent monoamine uptake has previously been reported directly following the COC-induced extracellular monoamine peak [52, 53]. Importantly, based on studies by others, vesicular monoamine release into the synaptic cleft is not only VMAT-2, but also Ca^2+^ and synapsin, dependent [54–56].

### Dopaminergic and serotonergic storage vesicles regulate the neurochemical response variation to COC

We found that COC increased the accumbal extracellular DA and 5-HT levels more strongly in HR than in LR rats. Although COC has previously been shown to inhibit the reuptake of monoamines by blocking plasmalemmal DA and 5-HT transporters, the finding that HR and LR rats did not differ in the cellular uptake of DA and 5-HT indicates that the observed individual differences in the extracellular accumbal DA and 5-HT response to COC cannot be attributed to individual differences in accumbal plasmalemmal DA and 5-HT transport. In fact, the results obtained in RES-treated HR and LR suggest that the individual-specific extracellular DA and 5-HT response to COC is mediated by release from storage vesicles. The observed larger dopaminergic and serotonergic storage capacity in the nucleus accumbens of HR compared to LR is in line with the fact that COC could still increase the extracellular accumbal DA and 5-HT levels in HR, but not in LR pre-treated with 1 mg/kg RES (see Supplemental Fig. S5). Accordingly, we conclude that COC leads to a stronger extracellular increase in DA and 5-HT in the nucleus accumbens of HR than of LR, because COC releases more DA and 5-HT from accumbal storage vesicles in HR than in LR (see Supplemental Fig. S5).

### Dopaminergic and serotonergic storage vesicles regulate behavioral response variation to COC

Our data not only support the generally accepted notion that accumbal DA regulates COC-induced walking [for review 57], but also suggest that COC-induced rearing is regulated by accumbal 5-HT. On the basis of the behavioral effects of RES, we hypothesize that the stronger rearing response to COC in HR than in LR is due to a larger COC-induced release of vesicular 5-HT in the nucleus accumbens of HR than of LR. Based on the neurochemical studies of Parsons and Justice [e.g. 58], it has frequently been suggested that the behavioral effects of accumbal 5-HT are due to 5-HT-receptor-mediated changes in the local release of DA [59, 60]. Our data not only indicate that accumbal 5-HT mediates rearing independently of changes in accumbal DA, but also that accumbal DA mediates walking behavior independently of changes in accumbal 5-HT. We hypothesize that the stronger walking response to COC in HR than in LR is due to a stronger COC-induced release of vesicular DA in the nucleus accumbens of HR than of LR.

### Monoaminergic storage vesicles control COC intake

The finding that 2 mg/kg RES reduced the ShA self-administration of COC, in addition to the finding that both 1 and 2 mg/kg RES reduced COC self-administration under LgA conditions, indicates that monoaminergic storage vesicles do not only control the COC-induced increase in the extracellular levels of accumbal DA and 5-HT, but also the voluntary intake of the psychostimulant. 1 mg/kg RES did not affect the ShA intake of COC, showing that the reduction in either COC intake, or in COC-induced walking and rearing, following RES-induced vesicle depletion does not result from motor impairment. The access-specific effect of 1 mg/kg RES also suggests that COC releases more monoamines from storage vesicles under LgA than under ShA conditions. This is in line with the observed faster recovery of the daily COC intake in ShA *versus* LgA rats treated with 2 mg/kg of the vesicle depleting agent.

### Storage vesicles control the individual-specific intake of COC

The positive correlation between locomotor response to novelty and COC self-administration is in line with the higher voluntary intake of this psychostimulant in HR than in LR [61–66]. The observed smaller reduction in the LgA COC intake in HR than in LR that were treated with 1 mg/kg of RES indicates that a larger vesicular DA and 5-HT storage capacity contributes not only to the stronger COC-induced increase in the extracellular levels of these monoamines, but also to the larger LgA intake of COC in HR than in LR. Not only the neurochemical and locomotor responses to COC, but also to high doses of amphetamine are dependent on the amount of monoamines inside storage vesicles [for refs 39, 57]. This indicates that the previously observed stronger locomotor response to amphetamine and the higher intake of this drug in HR than in LR [67–71] may also have resulted from a stronger amphetamine-induced release of DA and/or 5-HT from accumbal storage vesicles in HR than in LR. Intrestingly, the fact that 1 mg/kg RES did not affect the ShA intake of COC in both HR and LR, whereas this dose of the vesicle depleter was shown to strongly reduce the extracellular increase of accumbal monoamines in these rats, suggests that a mode of action different from an increase of extracellular DA and 5-HT in the nucleus accumbens may contribute to the rewarding effects of ShA exposure to cocaine. It remains to be investigated if this novel mode of action involves different neurotransmitters and/or changes in different brain regions.

### Storage vesicles may control the effects of COC in individuals lacking plasmalemmal transporters

Our finding that, apart from affecting plasmalemmal monoamine transporters, COC is able to increase extracellular monoamine levels may very well explain the previously reported unexpected intake of and behavioral response to COC in rodents lacking the plasmalemmal transporters [7, 8, 25, 26]. Such a monoamine-releasing action of COC may also explain why a reduction in plasmalemmal transporter gene expression does not eliminate the intake of COC by humans [22–24]. The transporter-independent mechanism of COC to increase the vesicular release of monoamines is still unknown. In this respect, it is interesting to note that COC stimulates sigma-1 receptors [for review 72], which in turn increases both 5-HT and DA release [73, 74].

## Conclusions

Altogether, our study shows that COC increases the extracellular concentrations of accumbal DA and 5-HT by depleting presynaptic storage vesicles, and that individual differences in the response to COC are mediated by individual differences in monoaminergic storage capacity. Therefore, storage vesicles may serve as an attractive novel drug target to develop inhibitors of psychostimulant intake. Our finding that RES inhibits the accumbal extracellular DA increase together with the generally accepted notion that accumbal DA mediates reward [1–4] suggests that RES reduced COC intake by interfering with the reinforcing properties of the psychostimulant. Given that extracellular 5-HT may also modulate reward [15, 20, 75, 76], future studies should focus not only on the manipulation of dopaminergic, but also of serotonergic storage vesicles as a potential therapy to inhibit the intake of COC.

## Acknowledgements

The authors wish to thank Benito Willemsen and Sandra van de Wiel for technical assistance and Dr. Dick Heeren for advice on statistical analysis of the data. MV and JH were supported by a joint program of the Netherlands Organization for Health Research and Development (ZonMW) and the USA National Institute for Drug Abuse (NIDA), project no. 31180005. MV is also supported by an ECNP research grant for young scientists and a NIDA INVEST Drug Abuse Research Fellowship. FF and LC were supported by grants from MIUR Progetto Eccellenza. The authors declare no conflict of interest.

