## supplemental material for "Action of cocaine involves depletion of dopaminergic and serotonergic storage vesicles"

| | |
|---|---|
| **Supplemental Materials and Methods** | **pages 2 – 10** |
| **Supplemental Fig. S1** | **page 11** |
| **Supplemental Fig. S2** | **page 12** |
| **Supplemental Fig. S3** | **page 13** |
| **Supplemental Fig. S4** | **page 14** |
| **Supplemental Fig. S5** | **page 15** |
| **Supplemental References** | **pages 16 - 17** |

### Supplemental Materials and Methods

#### Rats

All rats used were adult male Wistar rats of 180-220 g, bred at the Central Animal Facility of the Radboud University, Nijmegen, the Netherlands. They were housed in Macrolon cages (42 x 26 x 15 cm) under a fixed 12 h on / 12 h off light/dark cycle (lights on: 07.00 h) in a temperature-controlled room (21 ± 1.7 °C) and water and food were available *ad libitum*. Rats were handled 3 times per day for a period of 3 days before the day that they were either sacrificed or the microdialysis probe was inserted (see below). Experiments were performed in accordance with institutional, national and international laws and guidelines for animal care and welfare (see National Research Council (NRC) 2003 guidelines). Every effort was made to minimize the number of animals used and their suffering.

#### Experiment 1

#### Cocaine treatment and isolation of accumbal tissue

Unselected rats received a single injection of 45 mg/kg cocaine (COC) hydrochloride (Bufa, Spruyt Hillen, the Netherlands) or saline (volume: 1 ml/kg, i.p.) before being exposed to a test cage (diameter: 60 cm) with transparent walls (height: 40 cm) made of Plexiglas. A relatively high dose of COC was chosen, because it better resembles the high daily COC intake during the long access (LgA) self-administration experiments described below. At 20 min after this systemic injection, rats were sacrificed by decapitation, their brains were quickly removed, placed into a brain matrix, and sectioned at 3.0 mm intervals [1]. The first incision was made at the position where the two optic nerves fuse with the optic chiasm. From the identified slice, one punch of accumbal tissue was obtained from both hemispheres of the brain using a 1.22 mm i.d. needle [2]. The anterior commissure was used as a landmark to reliably punch out accumbal tissue (see Fig. 1A in the main document). The 20-min interval between drug injection and collection of the brain was chosen because at this time point the COC-induced increase in the extracellular levels of accumbal monoamines is known to be maximal [3].

**Vesicle isolation**

Purified accumbal vesicles were isolated by ultracentrifugation according to the protocol of Staal *et al.*, 2000 [4]. All centrifugation steps were performed in a Sorvall Micro ultracentrifuge (type: RC-M150GX, rotor: S120AT2) at 4°C. For each rat, punches of the left and right nucleus accumbens were pooled and homogenized in 1400 μl 0.32 M sucrose buffer (pH 7.3) containing 1 mM PMSF and 0.1 mg/ml soybean trypsin inhibitor. The homogenate was then centrifuged at 2000g for 10 min where after the resulting supernatant S1 was centrifuged at 10000g for 30 min. The resulting synaptosomal pellet P2 was re-suspended by swirling in 0.32 M sucrose buffer and subjected to an osmotic shock by the addition of ice-cold, distilled, deionized water. The osmolarity was restored by the immediate addition of 0.25 M HEPES and 1.0 M potassium tartrate buffer (pH 7.5). The osmotically shocked synaptosomes were centrifuged at 20000g for 20 min. Supernatant S3 was then centrifuged at 55000g for 60 min and $MgSO_4$ was added to supernatant S4 to bring the final magnesium concentration to 0.9 mM. The vesicular pellet P5 and its supernatant S5 resulted from the ultracentrifugation of supernatant S4 at 100000g for 45 min.

**Vesicular monoamine transporter 2 measurements**

The VMAT-2 Western blot procedures were adapted from Verheij *et al.*, 2008 [3]. Equal volumes (25 µl) of P5 and S5 were loaded on a 10% SDS-PAGE gel (total protein concentration of P5: 4 µg/µl) and electrotransferred to Protran nitrocellulose membranes (Schleicher and Schuell, USA). These membranes were exposed for 1h to blocking buffer (5% low-fat dry milk and 1% Tween-20 dissolved in PBS) where after they were incubated with rabbit anti VMAT-2 polyclonal antibody (1:500, AB1598P, Millipore, USA) overnight (at 4°C). After extensively rinsing with washing buffer (blocking buffer with 1% low-fat dry milk), blots were incubated with a secondary peroxidase-conjugated goat anti-rabbit antibody (1 : 5000, Nordic Immunology, the Netherlands) for 45 min (at room temperature), and subsequently washed. Peroxidase activity was immunodetected using Lumi-Light plus substrate (Roche Diagnostics, Germany). Hybridization signals were analyzed using Labworks 4.0 software (UVP bioimaging systems, UK). VMAT-2 levels were not normalized to tubulin, because the latter protein is not anymore present in vesicular fractions [5, 6].

**Determination of vesicular monoamine levels**

Vesicular pellet P5 was sonicated (Branson cell disruptor, USA) for 10 s in ice-cold tissue buffer (0.05 M sodium phosphate, 0.03 M citric acid buffer with 15% methanol (v/v), pH 2.5) and again centrifuged for 15 min at 100000g to remove membrane fragments. The final supernatant S6 was injected into a High Performance Liquid Chromatography (HPLC) system coupled to an electrochemical detector (ECD) for separation and quantification of DA and 5-HT (see section on microdialysis below for details). The vesicular monoamine levels of supernatant S6 were normalized for variation in protein loading using the total protein concentration of supernatant S1 (Bradford assay, Biorad, USA). The vesicular monoamine levels of each COC sample were calculated as a percentage of the mean of the vesicular monoamine levels of all saline samples measured on the same day to allow comparison across different centrifugation days.

**Walking and rearing**

In a previous study we showed that especially walking and rearing dominate behavior after the administration of COC [7]. The frequency of rearing (defined as events in which the front paw(s) are raised against the side wall(s) of the cage) and walking (defined as displacement of all 4 paws over a minimum distance of 1 cm for a period of at least 3 s) was scored by an observer blind to the rat type and its treatment, using a computer program (KEYS®) developed at our institute.

**Experiments 2-4**

**Selection of HR and LR**

High Responder (HR) and Low Responder (LR) to novelty rats [8-14] were selected according to procedures previously described in Cools *et al.*, 1990 [15]. Rats were placed on a square table of 160 x 160 cm that was made of black Perspex. This novel open field was 95 cm elevated above the floor, surrounded by a white neutral background (270 x 270 x 270 cm) and illuminated by white light of 170 Lux (see Supplemental Fig. S2). The selection of HR and LR rats to novelty is based on ambulation and habituation. Ambulation is defined as the overall distance traveled on the open field in a period of 30 min. Habituation time is defined as the duration of the period that starts as soon as the rat begins to explore the open field and ends as soon as ambulation stops for at least 90 s. Rats that habituated in less than 480 s and walked less than 4800 cm (in 30 min) were labeled LR whereas rats that habituated after 840 s and walked more than 6000 cm (in 30 min) were labeled HR.

**Determination of vesicular and extracellular DA and 5-HT levels in HR and LR**

**Vesicular and total DA and 5-HT measurements:** Nucleus accumbens punches of HR and LR rats were taken and fraction S6 (which consists of the re-suspended, sonicated and centrifuged vesicular pellet P5) was analyzed for vesicular DA and 5-HT levels according to the procedures described above. For quantification of the total levels of accumbal DA and 5-HT, nucleus accumbens punches of a new group of HR and LR rats were homogenized in phosphate-buffered saline (pH 7.3) containing 6 M Urea, 1% SDS, 1% β-mercaptoethanol, 1 mM PMSF and 0.1 mg/ml soybean trypsin inhibitor. The homogenate was sonicated for 30 s and centrifuged for 7 minutes at 20000 g, where after the supernatant was diluted in 0.1 M HCl (1:100) and immediately injected into the HPLC-ECD system (see below).

**Stereotactic surgery:** HR and LR rats were unilaterally implanted with a stainless steel guide cannula (length: 5.5 mm, outer diameter: 0.65 mm, inner diameter: 0.3 mm) directed at the right nucleus accumbens according to previously described procedures [16]. Under sodium pentobarbital anesthesia (60 mg/kg, volume: 1 ml/kg, i.p., pharmacy University of Utrecht, the Netherlands), rats were placed in a stereotaxic apparatus and the following coordinates were used: anterior: +10.6 mm (relative to the interaural line) and lateral: -1.5 mm (relative to the midline suture). The guide cannula was lowered 5.5 mm relative to the dura surface resulting in a vertical coordinate of +3.5 mm for the cannula tip. Finally, the cannula was angled 10$^{o}$ laterally to the right side. Coordinates were taken from Paxinos and Watson [17]. Screws and cement were used to fixate the cannula to the skull. The guide cannula contained an inner cannula to prevent infections and occlusions.

**Extracellular DA and 5-HT measurements:** After 7 days of recovery, a microdialysis probe (type A-I-8-02, outer diameter: 0.22 mm, 50000-molecular-weight cut-off, Eicom, USA) was inserted into the guide cannula. The tip of the dialysis probe protruded 2 mm below the distal end of the guide cannula. At 4 h following probe insertion, rats were injected with reserpine (RES, doses: 1 or 2 mg/kg, Daiichi, Japan) or its solvent (volume: 1 ml/kg, i.p.) and 24 h later accumbal dialysates were analyzed for DA and 5-HT using the below-mentioned HPLC + ECD system. As soon as the monoamine samples varied less than 10%, 5 baseline samples were taken. Immediately after the fifth baseline sample, rats were injected with 15 mg/kg cocaine (COC) hydrochloride or saline (volume: 1 ml/kg, i.p.) where after extracellular accumbal monoamine levels as well as walking and rearing were measured for an additional period of 120 min. RES-induced changes in the COC-induced increase of accumbal extracellular DA expressed as *percentage change of baseline levels* have previously been reported [3]. In the present study, we show changes in *absolute levels* of not only accumbal DA, but also accumbal 5-HT. At the end of the microdialysis experiments, rats were given an overdose of sodium-pentobarbital (250 mg/kg, i.p.) whereafter they were intracardially perfused with 60 ml 4% paraformaldehyde solution. Vibratome sections were cut and the location of the microdialysis probe was verified using the brain atlas of Paxinos and Watson [17].

**Microdialysis and HPLC + ECD system:** The inlet and outlet of the microdialysis probe were connected to a dual channel rotating swivel allowing the rat to move freely during the whole experiment. The COC dose of 15 mg/kg was chosen because this swivel system was not designed for the strong increase of stereotypic behavior (e.g. circling, head bobbing and gnawing) typically observed after higher doses of the psychostimulant. The probe was perfused at a rate of 2.0 µl/min with modified Ringer solution (147 mM NaCl, 4 mM KCl, 1.1 mM $CaCl_2.2H_2O$ and 1.1 mM $MgCl_2.6H_2O$, pH 7.4), and the outflow was automatically injected, once every 5 min, into a stand-alone HPLC+ECD system (type HTEC 500) of the Eicom company. 5-HT and DA were separated from the remaining neurotransmitters by means of reversed phase, ion-paring, liquid chromatography using an Eicompak PP-ODS column (particle size: 2 μm, 4.6 x 30 mm, Eicom) in combination with a mobile phase containing 1% of methanol (0.1 M phosphate buffer ($NaH_2PO_4.2H_2O$ : $Na_2HPO_4.12H_2O$, ratio 25:4), 2.0 mM sodium 1-decanesulphonate and 0.1 mM di-sodium EDTA, pH 6.0) at a flow rate of 500 $\mu$l/min (temp: 25 °C). The concentration of monoamines was measured by setting the working electrode of the electrochemical detector at + 400 mV against a silver/silver-chloride reference electrode. The HPLC-ECD unit was calibrated with a standard DA and 5-HT solution twice before each experiment. The detection limit of the electrochemical detector was about 30 fg (see www.eicom-usa.com).

**Determination of the reuptake of DA and 5-HT in HR and LR rats**

Accumbal punches of HR and LR rats were taken and synaptosomes were isolated as described above. The dose-dependent monoamine uptake into these synaptosomes was measured according to previous procedures [18, 19]. In short, the above-mentioned pellet P2 was re-suspended in 50 volumes of the original weight in Krebs-Henseleit-HEPES buffer. Tubes were pre-incubated for 5 min at 37 °C with 150 μl of this buffer containing 10-500 nM of [$^3$H]DA or 10-500 nM of [$^3$H]5-HT (25 Ci/mmol, Perkin Elmer, USA). Subsequently, 50 μl of the synaptosomal suspension was added and uptake was allowed for 5 min ([$^3$H]DA) or 20 min ([$^3$H]5-HT) at 37 °C. The uptake was terminated by adding ice-cold incubation buffer, followed by rapid filtration under vacuum through Whatman GF/B filters. The filters were washed with Krebs-Henseleit-HEPES buffer and the radioactivity trapped on the filters was counted by liquid scintillation. Total protein concentrations were determined in pellet P1 using a Bradford protein assay (Biorad, USA).

**Determination of COC self-administration in HR and LR**

**Surgery.** Rats tested for their locomotor response to novelty were implanted with a micro Renathane catheter (0.3 mm i.d. × 0.64 mm o.d.; MRE037, Braintec scientific Inc, USA) into the right external jugular vein according to previously reported procedures [20]. This aseptic surgery procedure was performed under 2-3% isoflurane anesthesia (TEVA pharmachemistry, the Netherlands). After surgery, rats were given analgesic (Flunixin, 2.5 mg/kg, s.c., Sigma-Aldrich, USA) and antibiotic (Cefazolin, 0.033 mg, i.v., Sagent Pharmaceuticals, USA) treatment for at least one week. The catheter was flushed twice daily with heparinized saline (30 USP/ml, Hospira, USA) during the entire experiment.

**Self-administration chambers.** COC self-administration was performed in standard operant chambers (28 x 26 x 20 cm, Med Associates Inc., USA) that were placed in ventilated, light- and sound-attenuating cubicles. The COC self-administration chambers were equipped with a swivel system allowing rats to move freely during self-administration sessions. COC was delivered by a 15 r.p.m. syringe pump (Razel Scientific Instruments, USA) and the start of a session was signaled by the presentation of 2 retractable levers into the self-administration chamber. Pressing the right lever was programmed to deliver cocaine (volume: 0.1 ml in 4s), whereas pressing the left lever had no programmed consequences. During drug administration, a stimulus light above the active lever was illuminated for 20 s indicating a timeout period when additional lever presses did not result in fluid delivery.

**COC self-administration measurements.** One week after surgery, HR and LR rats were trained to self-administer cocaine (COC) hydrochloride (0.5 mg/kg/infusion) under a fixed ratio 1 (FR1) schedule of reinforcement (one lever press resulted in one drug injection) for 1 h per day. Training was completed when the rats reached the criterium of 13 to 15 COC infusions for 3 consecutive days. After training, rats were divided into 2 groups matched by their number of infusions during the final training session. One group of rats continued to self-administer COC (0.5 mg/kg/infusion) in daily 1-h (short access) sessions, whereas the other group of rats self-administered the same COC dose in daily 6-h (long access) sessions [21, 22]. When the COC intake reached a stable level, rats were treated with RES (dose: 1 or 2 mg/kg) or its solvent (volume: 1 ml/kg, i.p.), where after they were exposed to their daily COC self-administration sessions for an additional period of 6 days.

**Analysis of the data**

Vesicular and total levels of 5-HT and DA were statistically analyzed using a one-way ANOVA. The relationship between these monoamine levels and behavior was evaluated by means of a Pearson's two-tailed correlation analysis (if HR and LR rats were used, the animals were pooled). The extracellular DA and 5-HT levels, the neuronal uptake of these monoamines, and COC self-administration were statistically analyzed using a three- or two-way ANOVA with the factors rat type, treatment and time (for repeated measures). Data are expressed as mean ± SEM and $p<0.05$ was considered statistically significant.

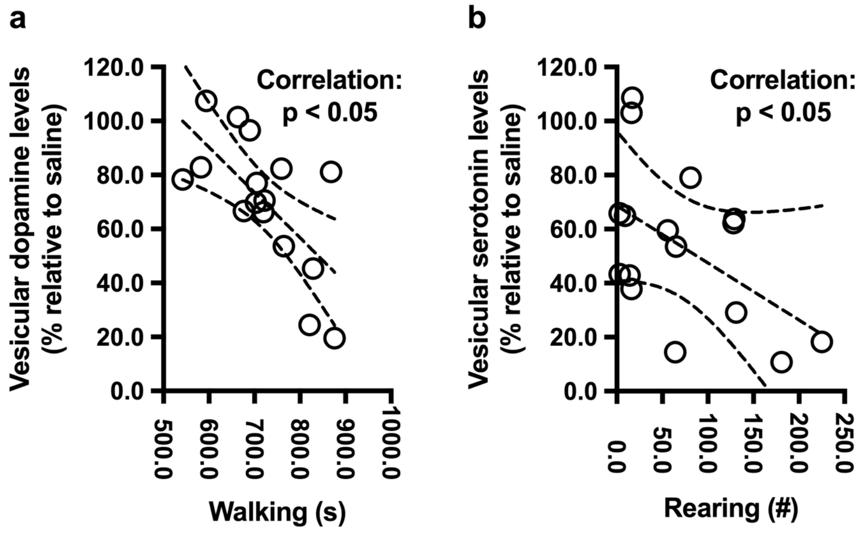


**Supplemental Fig. S1: Individual differences in cocaine-induced vesicle depletion.** Individual differences in the cocaine-(COC)-induced depletion of vesicular dopamine (DA) significantly correlated with individual differences in the COC-induced increase in walking, whereas individual differences in the COC-induced depletion of vesicular serotonin (5-HT) significantly correlated with individual differences the COC-induced increase in rearing (see also Fig. 1 of the main document).

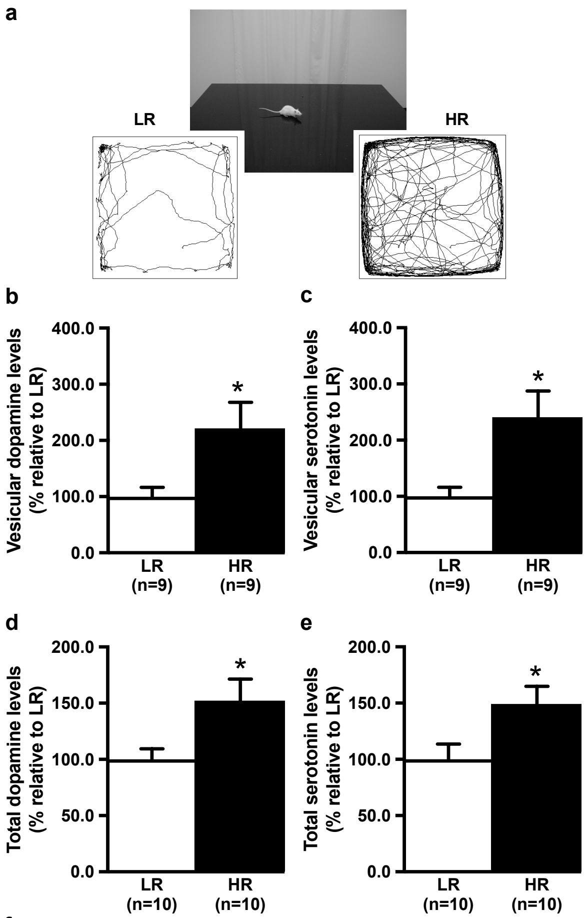


f

| | Distance on open field | Habituation time on open field |
|---|---|---|
| Vesicular dopamine levels | R = 0.534 p < 0.05 | R = 0.537 p < 0.05 |
| Vesicular serotonin levels | R = 0.572 p < 0.05 | R = 0.605 p < 0.01 |
| Total dopamine levels | R = 0.447 p < 0.05 | R = 0.454 p < 0.05 |
| Total serotonin levels | R = 0.519 p < 0.05 | R = 0.505 p <0.05 |

**Supplemental Fig. S2: Individual differences in vesicular monoamine levels.** A black open-field in a white neutral room was used to select Low Responder to novelty (LR) (left) and High Responder to novelty (HR) (right) rats (**a**). The vesicular and total levels of accumbal dopamine (DA) (**b/d**) and serotonin (5-HT) (**c/e**) were higher in HR than in LR, and these accumbal monoamine levels correlated significantly with both the distance travelled and the habituation time used to select these animals on the open-field (**f**). *Significant difference between HR and LR (one-way ANOVA: $p<0.05$). Correlation analysis was performed in the pooled group of HR and LR. Note: accumbal vesicular DA, but not 5-HT, levels have been taken from Verheij *et al.*, 2008 [3]. All data are expressed as mean ± SEM.

| | Basal dopamine levels | Basal serotonin levels |
|---|---|---|
| LR: reserpine solvent | 0.75 ± 0.18 | 0.12 ± 0.01 |
| LR: reserpine | 0.57 ± 0.09 | 0.07 ± 0.01 |
| HR: reserpine solvent | 0.82 ± 0.21 | 0.14 ± 0.01 |
| HR: reserpine | 0.32 ± 0.05 | 0.08 ± 0.01 |

**Supplemental Fig. S3: Effects of reserpine on basal monoamine levels.** Reserpine (RES) reduced baseline levels of extracellular accumbal dopamine (DA) and serotonin (5-HT) (two-way ANOVA: treatment effect: DA: $F_{(1,37)}=8.436$, $p<0.01$ and 5-HT: $F_{(1,37)}=25.969$, $p<0.001$) equally in High Responder to novelty (HR) and Low Responder to novelty (LR) rats (two-way ANOVA: rat type x treatment effect: DA and 5-HT: n.s.). Data represent the average of the 5 baseline samples (± SEM) shown in Fig.3 of the main document. Basal monoamine levels observed after 1 and 2 mg/kg reserpine were pooled (dose (x time) effect: n.s.).

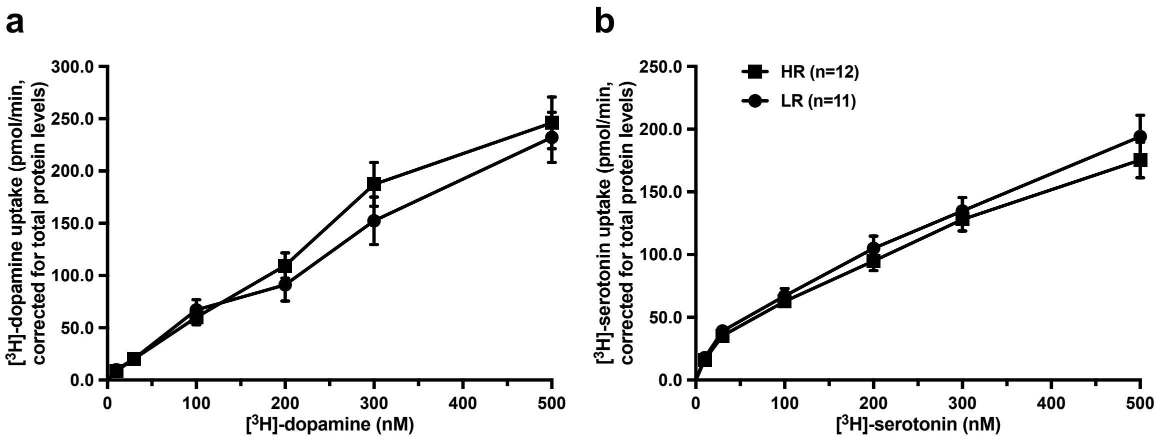


**Supplemental Fig. S4: No individual differences in monoamine reuptake.** The neuronal uptake of dopamine (DA) (**a**) and serotonin (5-HT) (**b**) in the nucleus accumbens of High Responder to novelty (HR) and Low Responder to novelty (LR) rats was concentration dependent, and did not differ between the two types of animals. All data are expressed as mean ± SEM.

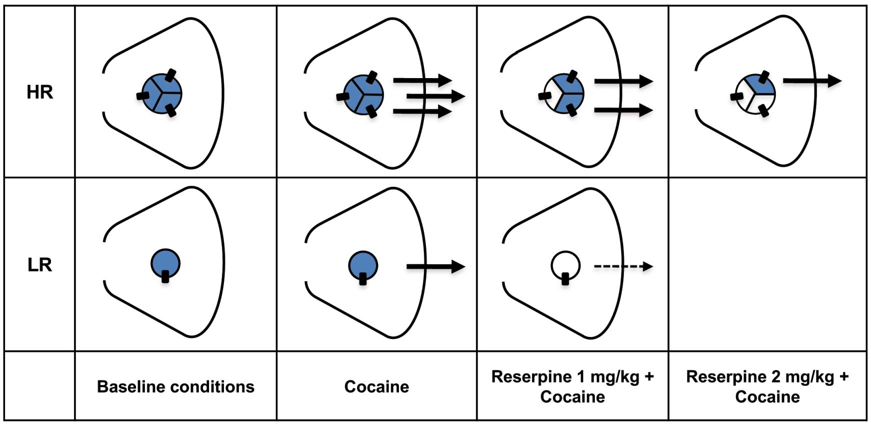


Suppl Figure S3

**Supplemental Fig. S5: Schematic representation of the results.** Circles ( ) represent monoaminergic storage vesicles inside the presynaptic dopaminergic or serotonergic neurons of the nucleus accumbens of High Responder to novelty (HR) and Low Responder to novelty (LR) rats. Black rectangles ( ) reflect vesicular monoamine transporters (number HR > number LR). Circle size reflects the monoaminergic storage capacity (capacity HR > capacity LR). Arrows represent the cocaine-(COC)-induced dopamine (DA) or serotonin (5-HT) release derived from vesicles (release HR > release LR). After reserpine (RES) treatment, vesicles become empty [i.e. blue circles become (partly) white], resulting in a reduced COC-induced release of vesicular DA or 5-HT into the synaptic cleft. Based on the neurochemical and behavioral effects reported in this study, we conclude that the COC-induced release of vesicular monoamines is larger in HR than in LR rats. Note: individual differences in vesicular monoamine transporter levels have been reported in Verheij *et al.*, 2008 [3].
